# A recurrent *de novo* splice site variant involving *DNM1* exon 10a causes developmental and epileptic encephalopathy through a dominant-negative mechanism

**DOI:** 10.1101/2022.06.02.492389

**Authors:** Shridhar Parthasarathy, Sarah M Ruggiero, Antoinette Gelot, Fernanda C Soardi, Bethânia F R Ribeiro, Douglas E V Pires, David B Ascher, Alain Schmitt, Caroline Rambaud, Hongbo M Xie, Laina Lusk, Olivia Wilmarth, Pamela Pojomovsky McDonnell, Olivia A Juarez, Alexandra N Grace, Julien Buratti, Cyril Mignot, Domitille Gras, Caroline Nava, Samuel R Pierce, Boris Keren, Benjamin C Kennedy, Sergio D J Pena, Ingo Helbig, Vishnu Anand Cuddapah

## Abstract

Heterozygous pathogenic variants in *DNM1* cause developmental and epileptic encephalopathy (DEE) due to a dominant-negative mechanism impeding vesicular fission. Thus far, pathogenic variants in *DNM1* have been studied using a canonical transcript that includes the alternatively spliced exon 10b. However, after performing RNA sequencing in thirty-nine pediatric brain samples, we find the primary transcript expressed in the brain includes the downstream exon 10a instead. Using this information, we evaluated genotype-phenotype correlations of variants affecting exon 10a and identified a cohort of eleven previously unreported individuals. Eight individuals harbor a recurrent *de novo* splice site variant, NG_029726.1(NM_001288739.1):c.1197-8G>A, which affects exon 10a and leads to DEE consistent with the classical *DNM1* phenotype. We find this splice site variant leads to disease through an unexpected dominant-negative mechanism. Functional testing reveals an in-frame upstream splice acceptor causing insertion of two amino acids predicted to impair oligomerization-dependent activity. This is supported by neuropathological samples showing accumulation of synaptic vesicles adherent to the plasma membrane consistent with impaired vesicular fission. Two additional individuals with missense variants affecting exon 10a, p.(Arg399Trp) and p.(Gly401Asp), had a similar DEE phenotype. In contrast, a single individual with a missense variant affecting exon 10b, p.(Pro405Leu), which is less expressed in the brain, had a correspondingly less severe presentation. Thus, we implicate variants affecting exon 10a as causing the severe DEE typically associated with *DNM1*-related disorders. We highlight the importance of considering relevant isoforms for disease-causing variants, as well as the possibility of splice site variants acting through a dominant-negative mechanism.

## Introduction

Developmental and epileptic encephalopathy (DEE) encompasses a group of disorders characterized by severe epilepsy refractory to medical treatment which affects children and results in delayed development, and both neurological and non-neurological comorbidities.^1–4^ Genetic factors are increasingly recognized as a major cause of DEEs, and more than 100 monogenic causes for epilepsies and neurodevelopmental disorders have been identified.^5–8^ Several common DEE-associated genes, such as *SCN1A* (MIM: 182389), *SCN2A* (MIM: 182390), *SCN8A* (MIM: 600702), and *STXBP1* (MIM: 602926), represent prime candidates for precision medicine approaches.^9–11^

Prominent among neurodevelopmental disorders and epilepsies with genetic etiologies are those due to variation in genes related to synaptic transmission. Several genes involved in synaptic vesicle fission and fusion, such as *DNM1* (MIM: 602377), *AP2M1* (MIM: 601024), *STXBP1, STX1B* (MIM: 601485), *VAMP2* (MIM: 185881), and *SNAP25* (MIM: 600322), have been implicated in neurodevelopmental disorders.^12–19^ In particular, *DNM1*-related disorders have a distinctive clinical presentation. Individuals with disease-causing variants in *DNM1* typically present with profound hypotonia from birth, cortical visual impairment, refractory infantile spasms with onset at 4—6 months of age that frequently evolve to Lennox-Gastaut syndrome with severe to profound developmental delay and intellectual disability.^17,19^

*DNM1* was first identified as a genetic etiology for epilepsy in 2014.^19^ *DNM1* encodes dynamin-1, a protein essential for clathrin-mediated endocytosis and synaptic vesicle recycling. Dynamin-1 oligomers perform the fission of clathrin-coated vesicles from the plasma membrane, thereby promoting vesicle-mediated neurotransmitter release at the synapse.^20,21^ Each dynamin-1 unit consists of a GTPase domain, middle domain, pleckstrin homology domain (PH), GTPase effector domain (GED), and proline rich domain (PRD).^20^ In mammals, this gene may lead to several isoforms distinguished chiefly by the alternative splicing of the tenth exon—some transcripts contain exon 10b and others the further downstream exon 10a.^22^

The mechanism of disease in *DNM1*-related disorders is primarily due to dominant-negative effects of mutant dynamin-1 secondary to missense variants.^17,23^ The dominant-negative mechanism of disease occurs when the mutant protein blocks the function of the wildtype protein product as well, causing the overall functional gene product to be reduced beyond what is expected from a single loss-of-function variant. In the case of *DNM1,* this is predicted to be due to formation of non-functional oligomers including mutant and wildtype dynamin-1 protein. Prior to the current study, it has been estimated that one-third of individuals have a single recurrent missense variant, NP_004399.2:p.(Arg237Trp); this and other recurrent variants, such as NP_004399.2:p.(Lys206Gln/Glu) and NP_004399.2:p.(Gly359Ala/Arg) are concentrated in the GTPase and middle domains.^17^

Haploinsufficiency, which occurs when only one wildtype gene copy is insufficient to preserve normal functioning of the protein product, is not a known mechanism of disease of *DNM1*; individuals with putative loss-of-function (pLoF) variants in *DNM1,* such as nonsense, frameshift, or splice site variants, have been identified in the healthy population.^24^

Nevertheless, there have been two previous reports of a likely disease-causing splicing variant, NG_029726.1(NM_001288739.1):c.1197-8G>A, which affects the alternately spliced exon 10a of this gene.^25,26^ The discovery of this possibly pathogenic splicing variant underscores the need to better understand the transcriptional profile of *DNM1,* which includes five known proteincoding transcripts in addition to 24 other predicted transcripts, mostly partial, in the Ensembl database.^27^

Here, we performed RNA sequencing in thirty-nine pediatric brain samples to understand the transcriptional complexity of *DNM1. We* find that the canonical *DNM1* transcript including the alternative exon 10b is not the predominant brain isoform. Instead, we identify transcripts containing exon 10a as the major cortical *DNM1* isoform. We extend this finding to understand genotype-phenotype correlations in 11 previously unreported individuals with disease-causing variants in *DNM1* affecting either exon 10a or exon 10b and find that disruption of exon 10a leads to the more severe, and more commonly reported, DEE phenotype.

## Materials and Methods

### Participant Recruitment

Informed consent for participation in this study was obtained from the parents of all subjects in agreement with the Declaration of Helsinki with approval by the Institutional Review Board (IRB) at respective institutions.

Six individuals included in the study were recruited at Children’s Hospital of Philadelphia (CHOP). Four individuals were referred by collaborating institutions: Children’s Hospital of San Antonio, USA (Individuals 5 and 6); Universidade Federal de Minas Gerais, Brazil (Individual 7); and Assistance Hôpitaux Publique de Paris, France (Individual 8). One individual was referred by a US clinical genetic testing laboratory.

### Genetic Analysis and Clinical Review

Prior to study inclusion, all individuals underwent diagnostic genetic testing, including gene panel (n=1) and whole-exome sequencing (WES, n=9). Gene panel testing in one individual was performed via whole-genome sequencing and targeted analysis of 1,362 genes at Hôpitaux Universitaires de Genève (Geneva, Switzerland). Whole-exome sequencing was performed in seven individuals by GeneDx with the IDT xGen Exome Research Panel v1.0 (Integrated DNA Technologies); in one individual by the Hospital for Sick Children (Toronto, Canada) using the Agilent SureSelect Focused Exome Kit (Agilent); in one individual at Hôpital de la Pitié Salpêtrière (Paris, France) using the SeqCap EZ MedExome Library kit (NimbleGen, Roche Sequencing); and in one individual by CHOP using the Agilent SureSelect Clinical Research Exome Kit (Agilent). Analysis, variant calling, and interpretation by ACMG criteria^28^ were performed using locally developed pipelines at each institution, and variants were confirmed using additional methods including Sanger sequencing for all probands.

De novo variants in all subjects mapped within or in proximity to exons 10a or 10b within the *DNM1* gene **(Figure 1A-B).** Each individual underwent a clinical data review of medical history information obtained from medical records and supplemented in nine individuals by parent interview conducted by a licensed genetic counselor (S.M.R., **Table 1**). Medical record review included developmental and seizure history, neurological findings, and morphological details. Available electroencephalogram (EEG) and brain imaging data were reviewed for all individuals. Seizure types were classified according to the International League Against Epilepsy (ILAE) classification criteria.^4^ An overall assessment of developmental status was made via chart review using the Gross Motor Functional Classification Scale (GMFCS) for gross motor function,^29^ either the Manual Abilities Classification Scale (MACS) for children over 4 years of age^30^ or the Mini-Manual Ability Classification System (mini-MACS) for children 1-4 years of age for hand use,^31^ and the Communication Function Classification System (CFCS) for communication skills.^32^ These ordinal scales were originally developed for children with cerebral palsy and are composed of five levels with scores of ‘I’ indicating higher levels of function while scores of ‘V’ representing severe limitations in function.

**Figure 1.**
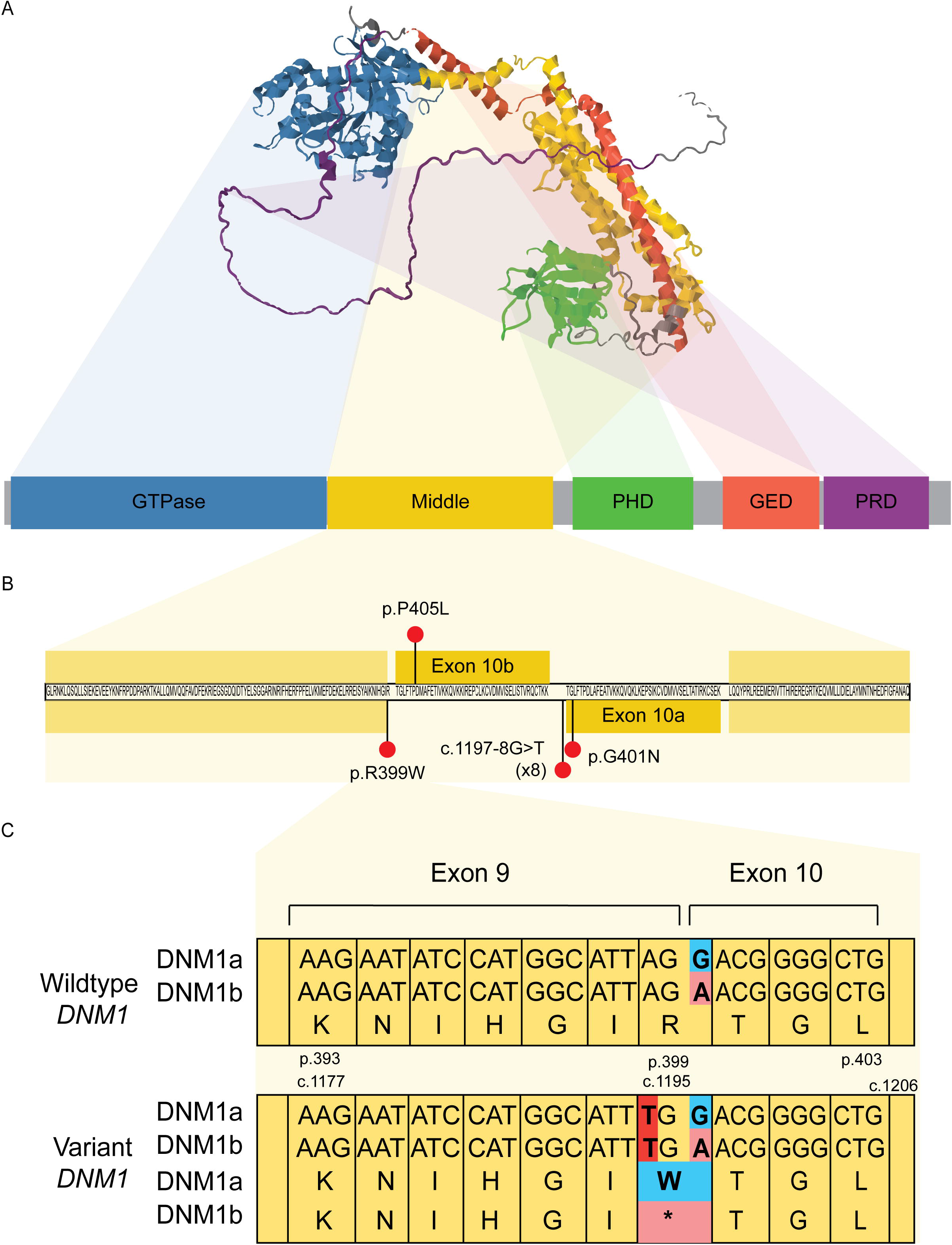
Structure and location of variants in *DNM1.* (A) Predicted 3D structure of dynamin-1 from AlphaFold and primary sequence of dynamin-1, highlighting distinct domains. (B) Subjects’ variants, including the recurrent variant NM_001288739.1:c.1197-8G>A, mapped to the dynamin-1 middle domain primary sequence, including alternative exons. The pathogenic variant-rich middle domain includes the alternatively spliced exon 10 of *DNM1*, with multiple disease-causing variants identified at positions uniquely affecting exon 10a. (C) Exon 10-dependent effects of the c.1195A>T variant. This variant is missense in DNM1a but nonsense in DNM1b; dominant-negative missense, not loss-of-function, variants are the established disease mechanism of *DNM1.*

### Assessment of Relative Transcript Expression

RNAseq was performed on brain tissue samples from 39 individuals from an independent cohort of individuals undergoing resective epilepsy surgery by Novogene Corporation Inc. using the NovaSeq 6000 PE150 platform (Illumina, Inc.). Raw short read files were quantified using Kallisto, version 0.45.0, against the GRCh38 human reference genome.^33^ Transcript isoform information was downloaded from Ensembl using the R package biomaRt.^34^ Transcript isoform abundance was estimated using output from Kallisto. Data were subsequently analyzed using R Statistical Framework. Transcript variants were filtered to exclude those less than 1500bp in length. The remainder were classified as containing exon 10a (“isoform DNM1a”) or 10b (“isoform DNM1b”). We assessed the correlation of proportional expression of each isoform with the age at sample collection using Pearson’s *r*, and we used Student’s *t* test to assess statistical significance of the ratio of DNM1a to DNM1b expression.

### Determination of Splicing Effects of Recurrent Intronic Variant

The MaxEnt and NNSPLICE algorithms were used within the Alamut software (Interactive Biosoftware, France) to predict the effect of the NM_001288739.1:c.1197-8G>A variant on *DNM1*.

The pCAS2 splicing reporter minigene assay was then used to experimentally determine the effect of this variant on the splicing pattern in *DNM1.* The *DNM1* amplification was performed with specific primers pair (DNM1_Forward 5’-GGGTCTTGTACGGAGCAGGG-3’ and DNM1_Reverse 5’-GAGTCAGATAGTAAGGGCAAGCAC-3’) using Pfu DNA polymerase (Promega). After the first amplification to select *DNM1* fragment of 761 bp, a nested-PCR was performed to construct the 494 bp minigene with the following primers: DNM1_Bam Forward 5’-CGG**<u>AT</u>**CCACGCTCACATTGTC-3’ and DNM1_Mlu Reverse 5’-GA**<u>C</u>**G**<u>C</u>**GTCATGAGTGCAAGTACC-3’. The restriction enzyme sites *BamHI* and *Mlu*I were inserted into the primer sequences to enable directional cloning. The minigene amplification product was inserted in pGEM T-easy vector (Promega) according to the manufacturer’s instructions. The construction was transformed into *E. coli* DH5α competent cells. After PCR amplification, clones with the correct insert size were isolated using the Wizard Plus SV Minipreps DNA Purification System (Promega) and sequenced to verify their identity. As the variant c.1197-8G>A was found in heterozygosis, minigene from patient wild-type sequence (normal allele) was used as control. pGEM wild-type and variant clones were digested with *Bam*HI and *Mlu*I restriction enzymes, respectively. The digested minigenes were purified with Wizard SV Gel and PCR Clean-UP System (Promega) and ligated into a *Bam*HI/*Mlu*I opened pCAS2 splicing reporter minigene vector (kindly provided by Dr. Alexandra Martins, Université de Rouen) (Gaildrat et al. 2010). The constructions were used to transform XL1B *E. Coli* competent cells (Stratagene), and colonies were selected to verify the presence of the *DNM1* fragment. Plasmid DNA was isolated using the Midprep kit (MN), and the sequence of the construct was verified by sequencing.

The pCAS2 wild-type and variant constructs were expressed by transient transfection into HEK293 cells. Approximately 5×10^5^ cells grown in each 60mm diameter tissue culture dish containing 4 mL Dulbecco’s modified Eagle’s medium-high glucose (Sigma-Aldrich) supplemented with 5 mM sodium bicarbonate (Cinética Química Ltda), penicillin (Sigma-Aldrich, 100 units/mL), streptomycin (Sigma-Aldrich, 0.1 mg/mL), amphotericin B (Sigma-Aldrich, 0.25 μg/mL), gentamicin (Schering-Plough, 60 mg/L), and 10% of fetal bovine serum (FBS) (Cripion Biotecnologia Ltda). After 24 hours, HEK293 cell cultures were transfected with 15 μg of the plasmid DNA using Lipofectamine 2000 (Invitrogen). Transfected cells were harvested 24 hours after transfection and total RNA was isolated using RNeasy mini kit (Qiagen) according to the manufacturer’s instructions, followed by DNase Amp Grade (Invitrogen) digestion. First-strand synthesis was performed using Superscript III kit (Invitrogen) according to the manufacturer’s recommendations. The cDNA was amplified by PCR using pCAS2 specific primers located in the flanking *SERPING1/C1NH* exons A and B.^35^ PCR products were analyzed by electrophoresis on a 1.5% agarose gel and the products were directly sequenced with *SERPING1/C1NH* exons A and B primers. Transfections were performed five times on different time points, no difference in the splicing results was observed between any of these independent assays.

### Structural Modeling of Splicing Variant

Solved three-dimensional structures of dynamin-1 were obtained by searching the RCSB Protein Data Bank (PDB). Several different X-ray crystal structures of dynamin-1 have been solved (PDB ID’s: 2X2E, 2X2F, 3ZYC and 3SNH), in addition to a medium resolution EM structure (PDB ID: 3ZYS) and an X-ray structure showing the tetrameric arrangements (PDB ID: 5A3F). The impact of the splicing variant was modeled on the tertiary structure of dynamin-1 based on these structures.^36^ Maestro was employed (Schrodinger, New York, NY), with wildtype and mutant structures minimized using the MMF94s forcefield in Sybyl-X 2.1.1 (Certara L.P., St Louis, MO), with the final structure having more than 95% of residues in the allowed region of a Ramachandran plot.

A comprehensive computational platform for analyzing mutation effects on protein structure function and interactions was employed to identify potential molecular mechanisms leading to the observed phenotype. For mutations located close to one of the protein-protein interfaces (<10Å), the mCSM-PPI2^37^ and mmCSM-PPI^38^ tools were employed to assess effects on oligomerization and binding affinity. Mutation effects on stability and dynamics were also assessed using the mCSM^39,40^ and DynaMut^41,42^ platforms. These provide a quantitative assessment of effects of mutations given as differences in Gibbs Free Energy (ΔΔG in Kcal/mol). Potential effects on GTP binding were also assessed using mCSM-lig.^43^

### Neuropathology Analysis

This study included one affected individual and one control individual obtained from the brain collection “Hôpitaux Universitaires de l’Est Parisien – Neuropathologie du développement” (Biobank identification number BB-0033-00082). For both cases studied, informed consent was obtained for autopsy of the brain and histological examination. After removal, brains were fixed with formalin for 5–12 weeks. Macroscopic analysis was performed allowing the selection and conditioning of samples (paraffin embedding, 7-micron slicing, hematein and periodic acid-Schiff (PAS) staining) of brain tissue for histological analysis. Immunohistochemistry was also conducted using anti-neurofilament 200 (Ready to use DAKO clone2F1 - reference: IS607) and anti-synaptophysin Ready to use clone DAKO-SYNAPTO - reference: IR660). Brains biopsies (globus pallidus, cervical spinal cord) were performed, and the samples were conditioned for electron microscopy as follows. Tissue was fixed in 3% glutaraldehyde in phosphate buffer, pH 7.4, for 1h, washed, post-fixed with 1% osmium tetroxide in 0.1 M phosphate buffer and then gradually dehydrated in 70, 90 and 100% ethanol. After 10 min in a 1:2 mixture of epoxy propane and epoxy resin and 10 mn in epon, coverslips were covered by upside down gelatin capsules filled with freshly prepared epoxy resin and polymerized at 60°C for 24 h. After heat separation from the coverslip, ultrathin sections of 90 nm were cut with an ultra-microtome (Reichert ultracut S), stained with uranyl acetate and Reynold’s lead and observed with a transmission electron microscope (JEOL 1011). Acquisition was performed with a Gatan ES1000W CCD camera.

## Results

### DNM1a is the predominant isoform expressed in the pediatric brain

*DNM1* encodes five known protein-coding transcripts in addition to 19 other predicted transcripts, mostly partial, in the Ensembl database.^27^ We determined relative expression of 24 transcripts, including two full-length transcripts containing exon 10a and three containing exon 10b, from RNAseq data in brain tissue samples following surgical resection in a pediatric cohort of 39 individuals without disease-causing *DNM1* variants **(Supplemental Table** 1). Thirteen individuals (33%) were female and 26 male (67%). 30 individuals (77%) had a diagnosis of focal epilepsy, and 9 (23%) had a diagnosis of epileptic encephalopathy. Intellectual disability or developmental delay was documented to be present in 15 individuals (38%) and absent in 24 (62%). Age of seizure onset ranged from 2 months to 13 years with a median of 4 years. The median age of sample collection was 8.6 years with a range of 14 months to 23 years.

Across this cohort with broad age range and diverse phenotypes, proportional expression of DNM1a, or transcripts containing exon 10a, was consistently higher than that of DNM1b, containing exon 10b, which includes the canonical transcript, with a mean ratio of 2:1 (Student’s *t* = 10.587, p < 0.001, **Figure 2A).** This expression pattern for the alternative tenth exons were not strongly correlated with individuals’ age (DNM1a: Pearson’s *r* = 0.22, p = 0.19; DNM1b: Pearson’s *r* = −0.46, p = 0.004; **Figure 2B),** reflecting consistency in the *DNM1* transcriptional landscape across early and adolescent development **(Figure 2C).** As such, DNM1a, which is distinct from the canonical *DNM1* sequence, is the primary cortical isoform with double the expression on average compared to DNM1b.

**Figure 2.**
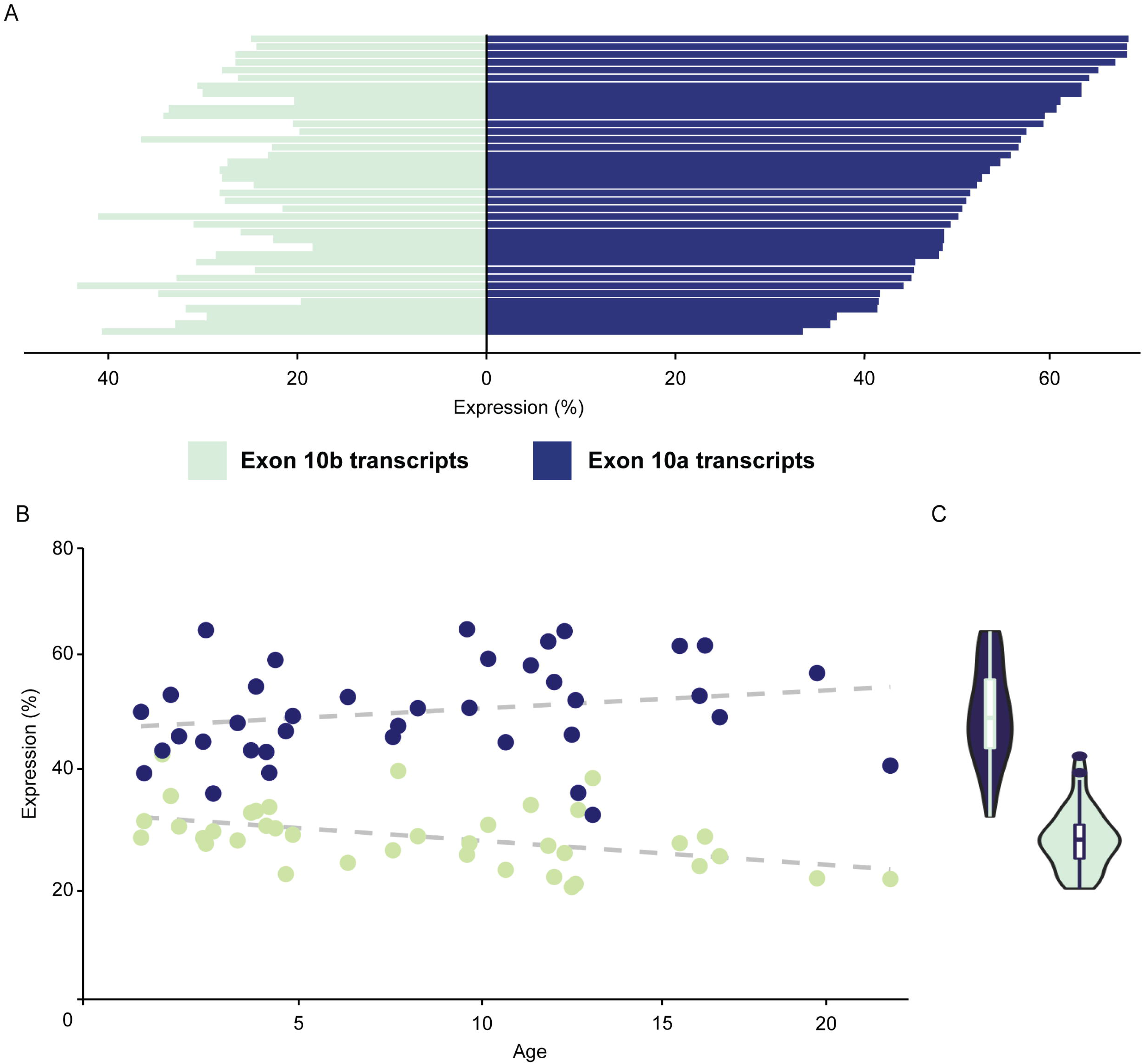
Expression profiles of *DNM1* transcripts from Ensembl in 39 pediatric brain tissue samples. (A) Individual transcript expression, highlighting identifiable transcripts within RefSeq. Bars represent individual samples arranged in order of increasing age at collection. (B) Proportional transcript expression for transcripts containing exon 10a and exon 10b vs age, showing age and expression are not strongly correlated. (C) Cumulative expression per individual of full-length *DNM1* transcripts containing exon 10a (blue) or exon 10b (green). DNM1a has markedly higher expression than DNM1b in the pediatric brain.

### Several disease-causing *de novo* variants in DNM1 affect exon 10a

Given that *DNM1* expression patterns strongly favor expression of exon 10a over exon 10b in the brain, we aimed to characterize clinical features of individuals with variants differentially affecting exon 10a or 10b **(Figure 1B).** Individuals 1-8 had a recurrent splice site variant,^25,26^ NG_029726.1(NM_001288739.1):c.1197-8G>A. This variant is directly upstream of the splice acceptor preceding exon 10a, and as such is expected to affect the splicing pattern of *DNM1* transcripts containing exon 10a but not 10b. Individual 9 had a missense variant, NM_001288739.1:c.1202G>A with the protein consequence NP_001275668.1:p.(Gly401Asp). This nucleotide substitution is within exon 10a and therefore results in an altered protein product from exon 10a-containing *DNM1* transcripts only. Individual 10 had a variant which was identified as nonsense, NM_004408.3:c.1195A>T with the protein consequence NP_001275668.1:p.(Arg399Ter). The affected codon includes the final two bases in exon 9 and the first base of exon 10a or 10b depending on the isoform. When assessed with a transcript containing exon 10b, as reported in clinical genetic testing, the nucleotide substitution changed this codon from CGA to TGA, a nonsense mutation inconsistent with known disease-causing variants. However, if exon 10a is considered instead, this substitution results in a change from CGA to TGG, with the protein consequence NP_004399.2:p.(Arg399Trp) **(Figure 1C).** This is a missense variant in the middle domain, which is known to be enriched for disease-causing variants with dominant-negative effects.^17^ Individual 11 had a missense variant, NM_004408.3:c.1214C>T with the protein consequence NP_004399.2:p.(Pro405Leu). This is a nucleotide substitution within exon 10b, leading to a missense variant in isoforms including this exon but not in those including exon 10a.

### A recurrent splice variant causes an in-frame insertion disrupting *DNM1* oligomerization

*DNM1*-related disorders are known to cause DEE through a dominant-negative disease mechanism.^17,23^ However, although the phenotypes of Individuals 1-8 mirrored the previously described DEE phenotype, splice site variants are typically assumed to be putative loss-of-function variants, which do not have dominant-negative effects. Therefore, we investigated the functional consequences of this splice site variant. First, we evaluated the effects of on dynamin-1 monomers. We examined this variant with multiple splice site prediction algorithms and found that the NM_001288739.1:c.1197-8G>A variant is predicted to create a new cryptic acceptor splice site 6 bps upstream of the canonical site of exon 10 in the stalk of the middle domain. The predicted change at the canonical site is −80.9% according to the MaxEnt algorithm and −44.5% according to the NNSPLICE algorithm.

To verify the validity of these *in silico* predictions, we also investigated the splicing variant experimentally with the pCAS2 splicing reporter minigene assay. By sequencing amplified cDNA following transfection of wildtype and mutant *DNM1* into HEK293 cells, we confirmed that the c.1197-8G>A variant indeed created a new splice site upstream of the canonical splice site, leading to the insertion of two amino acid residues, a cysteine and an arginine (CR) in the beginning of exon 10, between Arg399 and Thr400 in the stalk region of the middle domain **(Figure 3A-B).**

**Figure 3.**
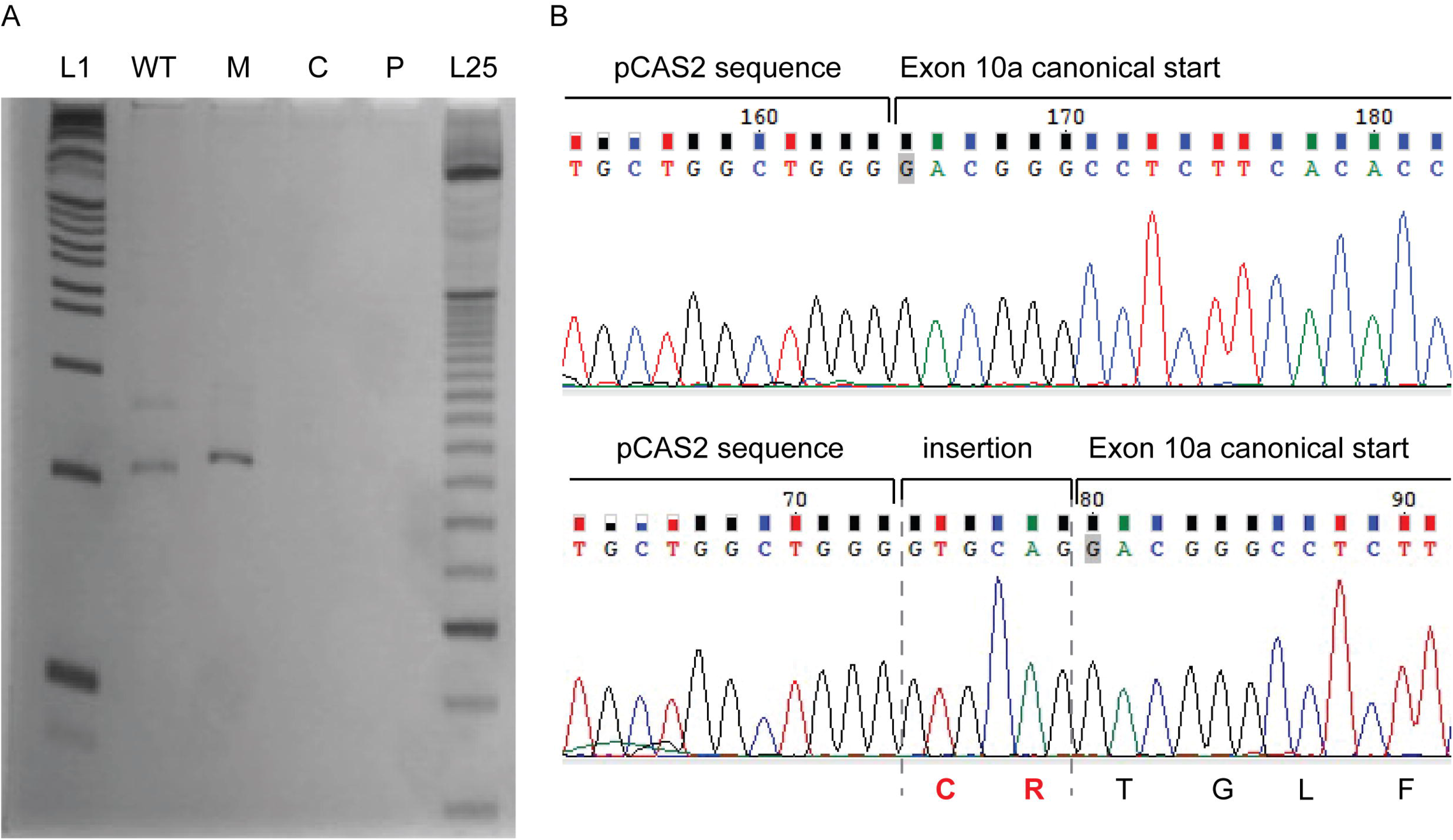
Splicing assay assessing consequence of NM_001288739.1:c.1197-8G>A variant. (A) cDNAs PCR products. L1 = 1Kb plus DNA ladder (Invitrogen), WT = wild-type amplification, M = mutant amplification, C = amplification from HEK293 cells without minigene transfection, P = PCR control without cDNA, L25 = 25 bp ladder (Invitrogen). (B) Sanger sequencing of Wild-type and Mutant PCR products. In red, the insertion of “GTGCAG” from 5’ of intron 9 to the mutant transcript, which results in a mutant protein with a cysteine and an arginine (CR) in frame insertion after Arg399 residue.

To discern the impact of this two-amino acid insertion on the tertiary and quaternary structures of dynamin-1, we modeled this variant based on established structures of the protein. Using several previously described computational tools,^36-43^ we found that in the protomeric state the insertions were on a solvent exposed loop, with minimal effect to monomer stability (ΔΔG = +0.2 Kcal/mol). However, the insertion was located at one of the interfaces within the dynamin-1 tetramer, and two extra residues would sterically hinder the formation of the active tetramer. The tetramer is formed by a pair of parallel protomers. When a mutant protomer containing the CR insertion formed part of the complex, the corresponding parallel protomer **(Figure 4A-B;** *blue)* could no longer bind due to steric hindrance **(Figure 4A-B).** Consequently, the insertion results in a mutant protein with normal interactions through the GTPase domain and with one anti-parallel protomer but disrupted interactions with one parallel protomer and the remaining anti-parellel protomer **(Figure 4C-D),** forming an oligomeric complex without the capacity for oligomerization-induced GTPase activation. By thus sequestering wildtype proteins needed to form the active complexes, the NM_001288739.1:c.1197-8G>A variant inactivates the enzyme without affecting monomeric structure. As such, a subset of wildtype dynamin-1 would be prevented from exhibiting normal function, reflecting a dominant-negative mechanism that is known to cause severe disease.

**Figure 4.**
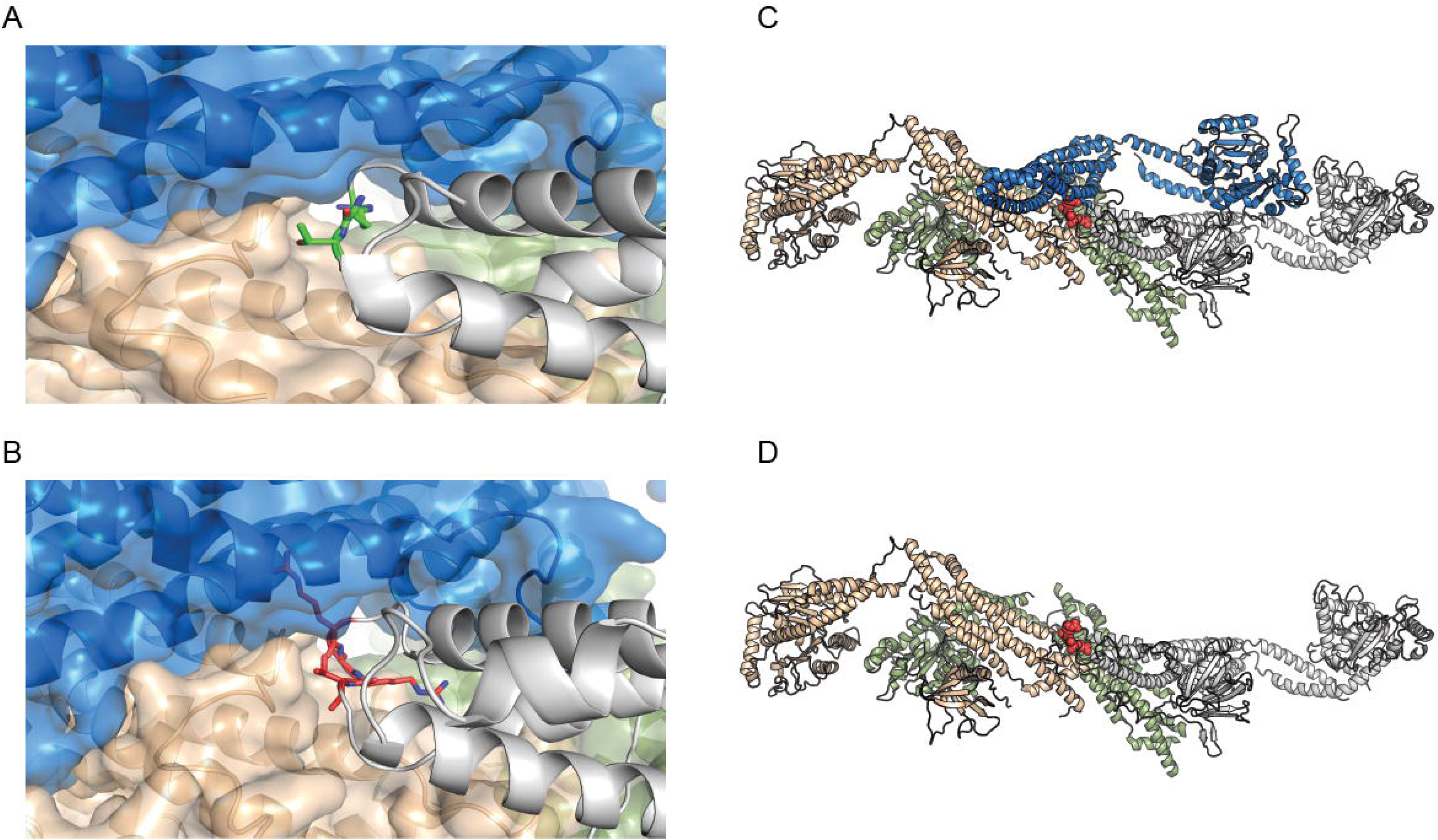
Structural modelling of the effects of the NM_001288739.1:c.1197-8G>A variant. (A) Mutation site in dynamin-1 corresponding to NM_001288739.1:c.1197-8G>A. Green = two residues neighboring the CR insertion. (B) Model of the CR insertion (pink). Given the lack of space to accommodate, the new residues steric clashes would be created, disrupting the interactions to the neighboring protomers. (C-D) Tetrameric organization of DNM1 (PDB ID: 5A3F). The tetramer is formed by two sets of parallel protomers facing in the same direction. Grey and green; and blue and orange. When a mutant protomer containing the CR insertion (bottom- grey chain) forms part of the complex, the corresponding parallel protomer (blue) can no longer bind due to steric hindrance.

### Neuropathology reveals vesicular accumulation in an individual with NM_001288739.1:c.H97-8G>A

Having established that the recurrent variant NM_001288739.1:c.1197-8G>A is predicted to cause dominant-negative disease, we next sought to assess its effects on native vesicular function. In the presynaptic terminal, neurotransmitter release relies on vesicles fusing with the membrane and subsequently being recycled via endocytosis. Dynamin-1 is known to perform vesicle fission, or the process of pinching synaptic vesicles off the plasma membrane. Thus, we hypothesized that this variant would lead to evidence of impaired synaptic vesicle fission. In one deceased individual (Individual 8), neuropathology analysis permitted detailed exploration of the effects of this recurrent splice variant on the synapse. This individual had significantly reduced brain weight (902g compared to an expected 1150g), with diffuse atrophy of cortical grey and white matter as well as central reduction of white matter **(Figure 5A-B).** Infratentorial structures appeared macroscopically preserved.

**Figure 5.**
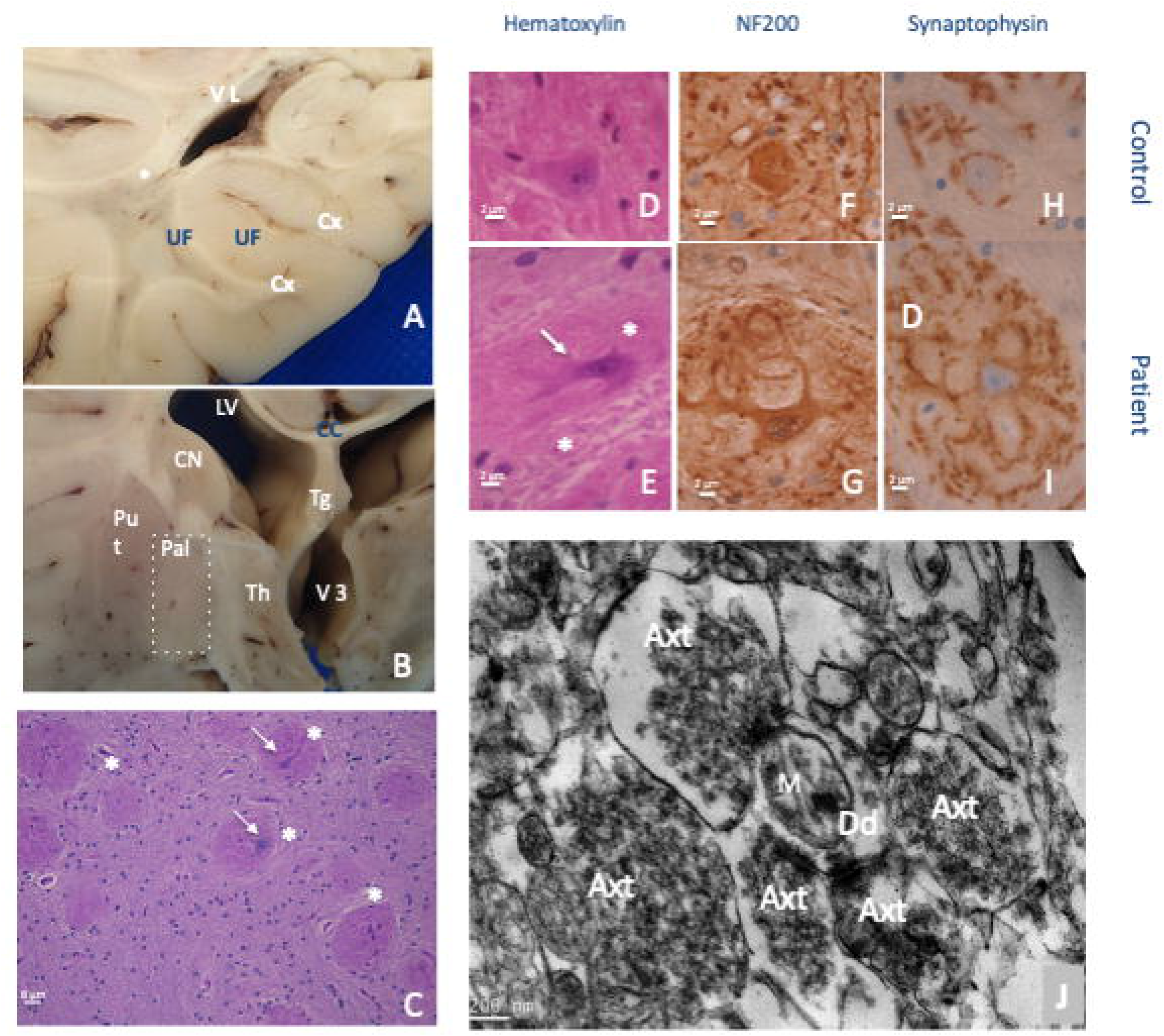
Neuropathological hallmarks of the NM_001288739.1:c.1197-8G>A variant. (A,B) Brain macroscopic aspect. (A) White matter displayed severe atrophy with lateral ventriculomegaly, spindly gyri and extremely thin corpus callosum. A central myelin discoloration was observed consisting of a myelin defect sparing U fiber. (B) Among grey structures, thalamus was severely atrophic associated with a third ventricle distention. Cortical thickness seemed preserved but was difficult to estimate considering the associated severe white matter atrophy. CC = corpus callosum, Cx = cortex, LV = lateral ventricle, Put = putamen, Pal = pallidum, Th = thalamus, Tg = trigone, UF = U fibers, V3 = third ventricles, * = deep white matter. (C-J) Pallidal synaptic dysplasia. (C) Pallidal glomeruli appeared as rounded formations containing intermingled eosinophilic neurites. (D-I) After immunohistochemistry and control matching, these glomeruli consisted of exuberant neurites proliferation (NF200(+)) that were covered by diffuse synaptic areas (synaptophysin). D,F,H are age-matched controls; F,G are immunohistochemistry with anti-neurofilament 200; H,I are with anti-synaptophysin; initial magnification x100. (J) Brain biopsy at pallidum level after electron microscopy conditioning. Initial magnification: x7500. Five hyperplastic synaptic terminations surrounding one receptive dendrite. Synaptic vesicles appeared abnormally numerous and packed.

At the histological level, we identified three types of lesions. (1) In grey structures, we observed decreased neuronal density, which was severe in the thalamus and pyramidal layers of the cortex and less marked in the striatum and globus pallidus. (2) In white matter, we observed that the leukodystrophy was related to a decrease in the density of axons whose myelin was preserved, i.e., no naked axons or hypercellularity. (3) We found a diffuse synaptic dystrophy, particularly marked at histological level in the internal globus pallidus, where the soma of the pyramidal neurons were embedded by glomerules consisting of a lace of exuberant dendrites displaying hyperplastic synaptic zones **(Figure 5C-I, Figure 6G).** Similarly, in the cerebellar cortex, synaptic areas were hypertrophic in the internal granular layer and increased in size and number around the soma of Purkinje cells **(Figure 6A-D).** At the ultrastructural level, hypertrophic synaptic terminals **(Figure 5J)** were filled by vesicles were irregular in size and shape, sometimes displaying a tubular shape **(Figure 6E),** and densely packed against the cell membrane **(Figure 6F);** these features were consistent with failed vesicular fission. In addition, we observed signs of intra-axonal trafficking defect, with accumulation of organelles such as vesicular bodies or mitochondria **(Figure 6H-J),** the latter displaying abnormalities in size, shape, and crest organization. Comparable features, i.e., excessive visibility of synaptic zones and axonal trafficking defect, were also observed in the peripheral nervous system by ultrastructural analysis of the skin biopsy samples **(Figure 6K-M).** Taken together, these findings reveal marked synaptic structural abnormalities with an accumulation of vesicles at the plasma membrane, consistent with disruption of vesicular fission due to mutant dynamin-1.

**Figure 6.**
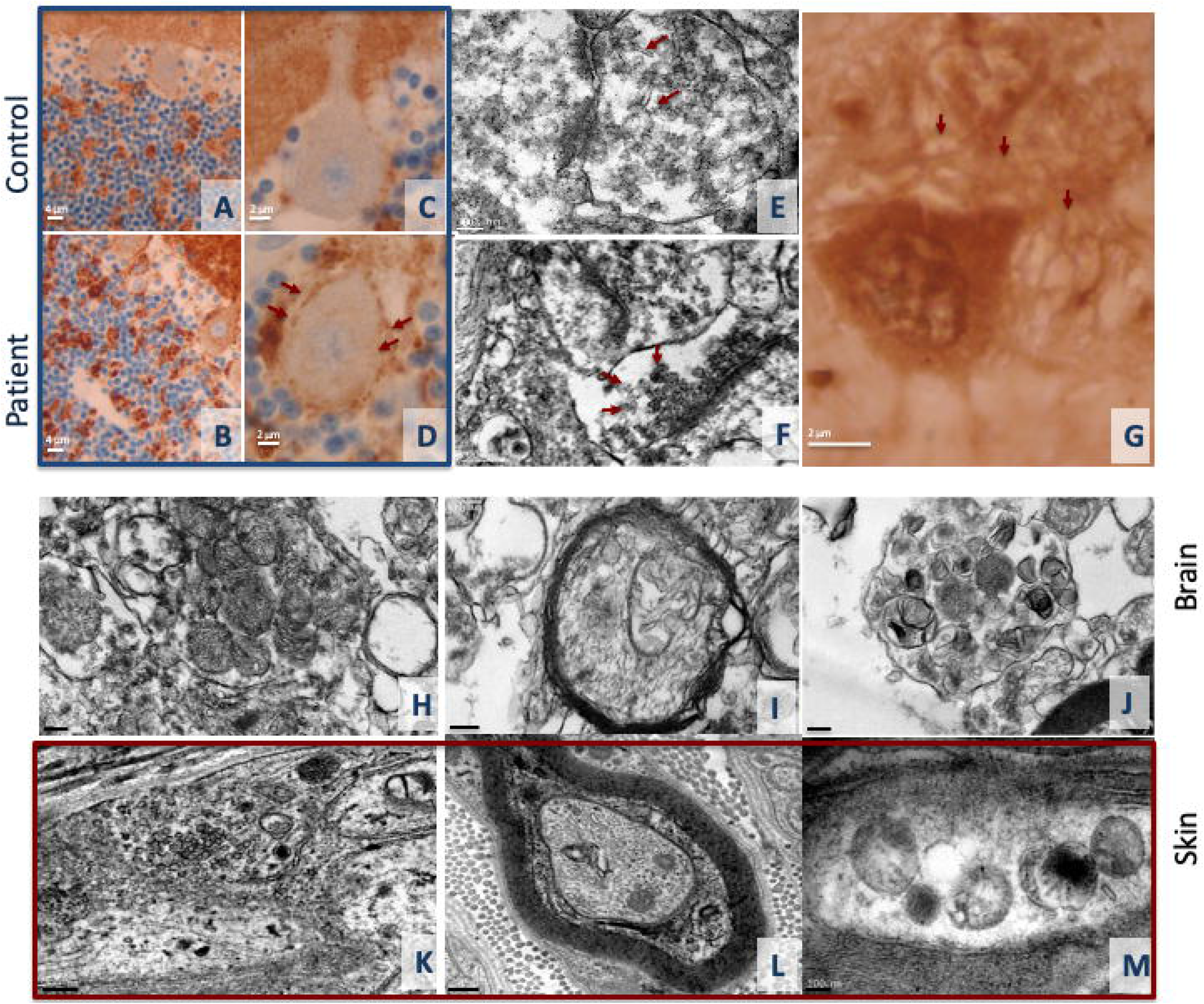
Synaptic dysplasia expression in brain and skin biopsy. (A-D) Cerebellar cortex revealed an excess of synapses as disorganization of granular glomeruli; A,B with initial magnification x40 and C,D with x100; A,C are age-matched controls. (E-H) Synaptic dysplasia consisted of (G) dendrite proliferation evident in pallidal glomeruli; (E-F) excess synaptic vesicles in thalamus with abnormal shape, irregular size, densely packed and adherent to synaptic membrane; (H-J) excess mitochondria in cervical spinal cord axon with abnormal appearance, with crests irregular in size, rounded and disorganized. E with initial magnification x15000, F with x12000, G with x100, H with x7500, I with x12000, J with x7500. (K-M) In skin biopsy, most of the observed axons contained heterogenous membranous vesicles that are not pathognomonic features but suggest altered trafficking. However, synaptic areas (axonal zone filled by synaptic vesicles), which are typically difficult to find in skin biopsy, were easily observed in this individual. K with initial magnification x12000, L with x10000, M with x15000.

### Clinical features in individuals with exon 10a variants reflect the classical *DNM1* phenotype

Ten individuals in this cohort had *de novo* variants in *DNM1* affecting exon 10a. All 10 of these individuals presented with a *DNM1*-related disorder consistent with the previously described DEE phenotype.^17^ Individuals 1-10 presented with early-infantile onset epileptic encephalopathies, including highly abnormal EEGs with or without clinical seizures (n=10). Two individuals passed away, one in infancy and one in young childhood, due to complex feeding difficulties and severe failure to thrive. Individuals presented with either profound (n=9) or severe (n=1) global developmental delay. All children were classified as Level V on the GMFCS, MACS/mini-MACS, and CFCS with the exception of Individual 4, indicating severe functional limitations in gross motor, fine motor, and communication skills **(Table 1)**. The oldest child included in the cohort, Individual 4, a 7-year-old, was able to communicate verbally with a few individual words and using a picture board (CFCS Level IV), and he was able to stand, but not walk, with assistance (GMFCS Level V). Developmental and seizure outcomes for the remaining (n=9) individuals are limited by their young age, but the remaining individuals had achieved neither head control nor the ability to communicate with spoken language by the time of the last clinical evaluation. Eye contact and visual tracking were reduced due to cortical visual impairment (n=9). Epilepsy presentations in all individuals were characterized by early-infantile onset seizures and abnormal EEG findings in 8/8 individuals **(Figure 7C),** or onset of infantile spasms with hypsarrhythmia at 6 months of age (n=2). One individual had highly abnormal EEGs significant for polymorphic slow waves seen diffusely as well as multifocal epileptiform discharges, at times concerning for hypsarrhythmia. She had not experienced clinical or subclinical seizures. Seizures remained refractory for most individuals (n=9); Individual 4, however, had initially very refractory epilepsy but achieved excellent seizure control with combination therapy of medicinal marijuana extract, valproic acid, and lamotrigine. Notable MRI features **(Figure 7A-B)** included decreased cerebral volume over time (3/7 individuals) and decreased myelination (2/7).

**Figure 7.**
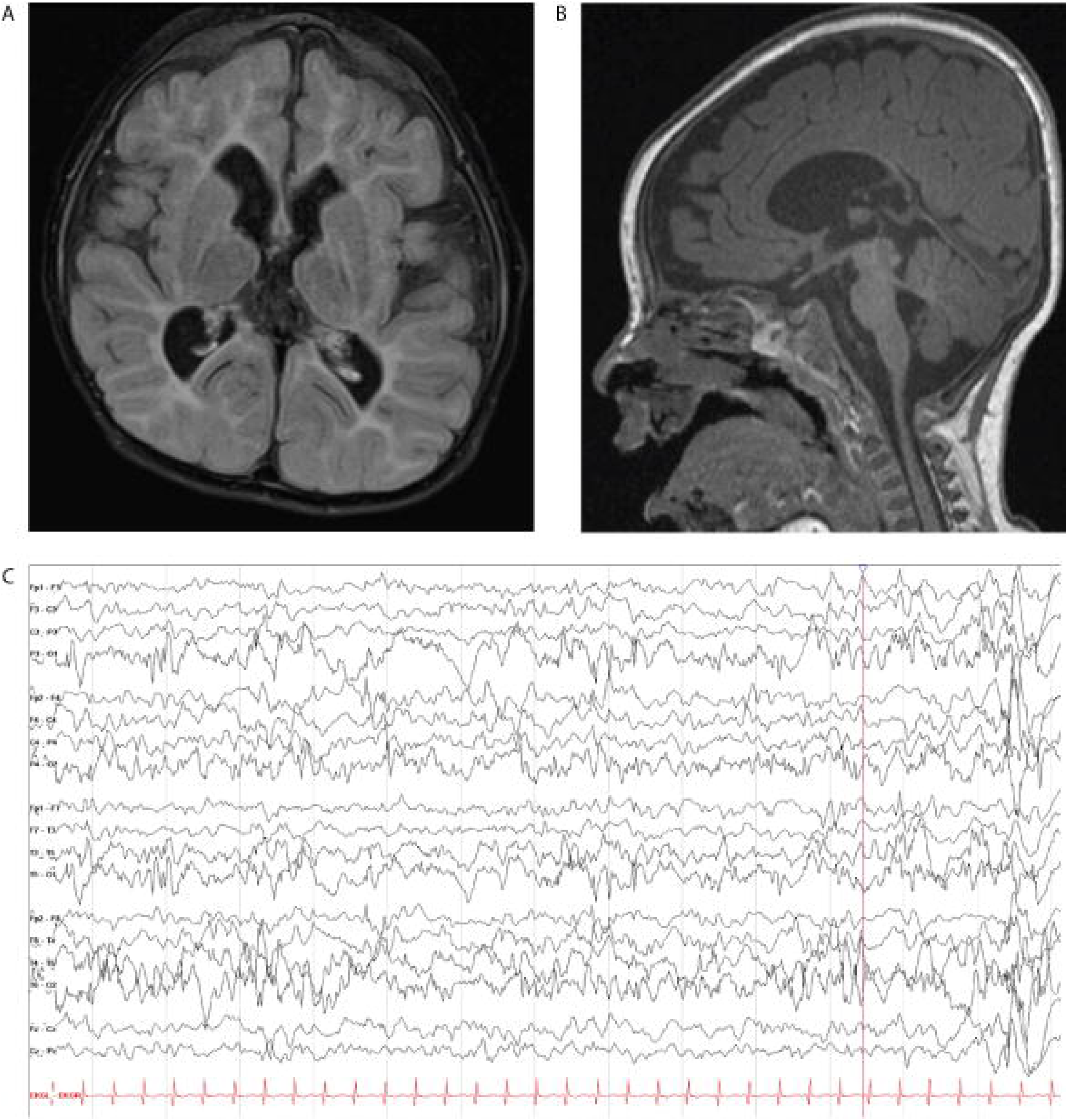
Imaging and electroencephalography from Individual 1 with the recurrent splice variant NM_001288739.1:c.1197-8G>A. (A-B) MRI showing markedly decreased cerebral volume, with deficiency of cerebral white matter. (A) Coronal FLAIR. Slightly dysmorphic lateral ventricles are enlarged, with prominent frontal horns and mild prominence of the temporal horn. (B) Sagittal T1. Small brainstem and small corpus callosum (C) Abnormal EEG from the same individual with poorly organized, diffusely slow background; nearly continuous focal slowing in the left occipital region; and multi-focal epileptiform discharges.

### One individual with a variant exclusively affecting exon 10b has atypically mild features

A 5-year-old girl presented with a *de novo* variant in *DNM1,* NM_004408.3:c.1214C>T, which is located within exon 10b. This child presented to care when she was first noted to have global developmental delays in infancy and diffuse hypotonia was diagnosed at this time. She achieved walking independently at 18 months of age, and at age 4 she was able to use verbally communicate with multiple words short sentences. When she was evaluated at age 4, was classified as Level I on the GMFCS and CMFCS, which indicates that she able to walk and climb stairs independently, as well communicate easily with both familiar and unfamiliar people. However, due to her hypotonia and cognitive delays, she had decreased standing balance during gross motor play activities and used supramalleolar orthotics to increase ankle stability. She had difficulty following one step verbal instructions and had behavioral problems with decreased safety awareness. She had distinct difficulties with visual-motor skills resulting in a MACS classification of Level II indicating she can manipulate objects with decreased speed and quality of movement (Table 1). An MRI obtained following *DNM1* diagnosis was normal, and her EEG was significant only for increased beta frequencies. At the time of the last clinical evaluation at age 4, she never had clinical seizures.

**Table 1.**
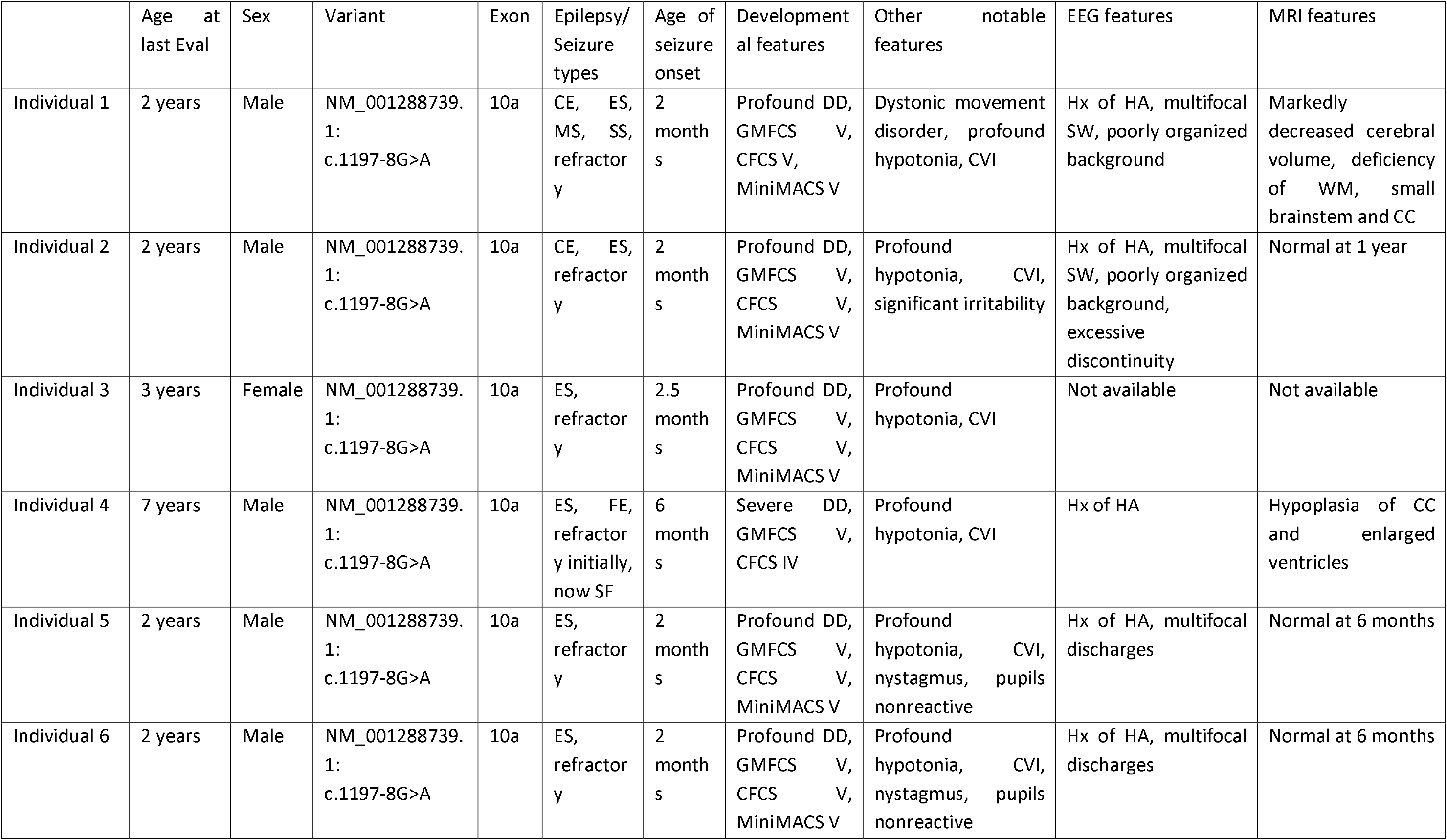
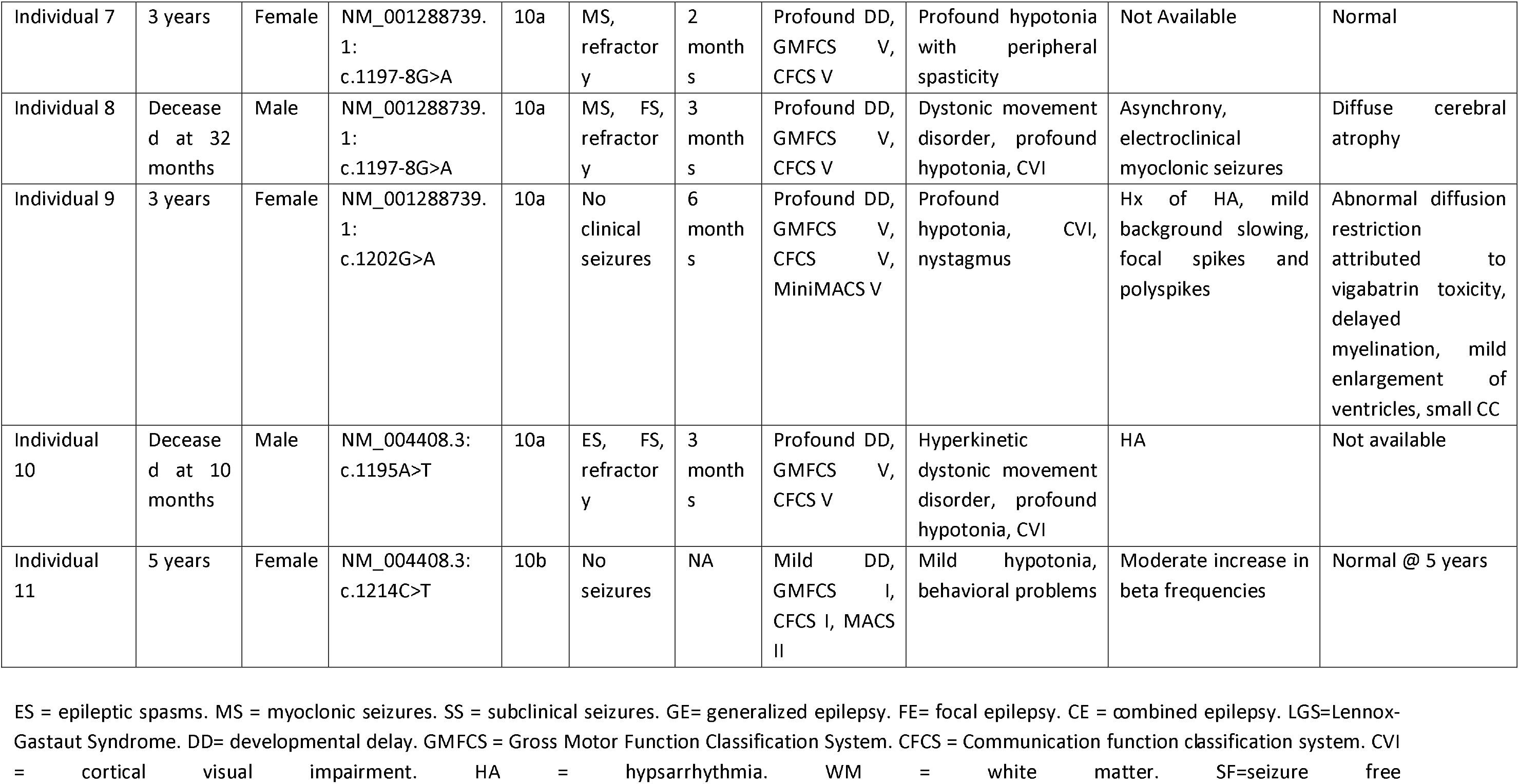
Clinical and genetic features in 11 individuals with *DNM1*-related disorders.

|  | Age at last Eval | Sex | Variant | Exon | Epilepsy/ Seizure types | Age of seizure onset | Developmental features | Other notable features | EEG features | MRI features |
| --- | --- | --- | --- | --- | --- | --- | --- | --- | --- | --- |
| Individual 1 | 2 years | Male | NM_001288739.1:<br>c.1197-8G>A | 10a | CE, ES, MS, SS, refractory | 2 months | Profound DD, GMFCS V, CFCS V, MiniMACS V | Dystonic movement disorder, profound hypotonia, CVI | Hx of HA, multifocal SW, poorly organized background | Markedly decreased cerebral volume, deficiency of WM, small brainstem and CC |
| Individual 2 | 2 years | Male | NM_001288739.1:<br>c.1197-8G>A | 10a | CE, ES, refractory | 2 months | Profound DD, GMFCS V, CFCS V, MiniMACS V | Profound hypotonia, CVI, significant irritability | Hx of HA, multifocal SW, poorly organized background, excessive discontinuity | Normal at 1 year |
| Individual 3 | 3 years | Female | NM_001288739.1:<br>c.1197-8G>A | 10a | ES, refractory | 2.5 months | Profound DD, GMFCS V, CFCS V, MiniMACS V | Profound hypotonia, CVI | Not available | Not available |
| Individual 4 | 7 years | Male | NM_001288739.1:<br>c.1197-8G>A | 10a | ES, FE, refractory initially, now SF | 6 months | Severe DD, GMFCS V, CFCS IV | Profound hypotonia, CVI | Hx of HA | Hypoplasia of CC and enlarged ventricles |
| Individual 5 | 2 years | Male | NM_001288739.1:<br>c.1197-8G>A | 10a | ES, refractory | 2 months | Profound DD, GMFCS V, CFCS V, MiniMACS V | Profound hypotonia, CVI, nystagmus, pupils nonreactive | Hx of HA, multifocal discharges | Normal at 6 months |
| Individual 6 | 2 years | Male | NM_001288739.1:<br>c.1197-8G>A | 10a | ES, refractory | 2 months | Profound DD, GMFCS V, CFCS V, MiniMACS V | Profound hypotonia, CVI, nystagmus, pupils nonreactive | Hx of HA, multifocal discharges | Normal at 6 months |
| Individual 7 | 3 years | Female | NM_001288739.1:<br>c.1197-8G>A | 10a | MS,<br>refractor<br>y | 2<br>month<br>s | Profound DD,<br>GMFCS V,<br>CFCS V | Profound hypotonia<br>with peripheral<br>spasticity | Not Available | Normal |
| Individual 8 | Deceased at 32 months | Male | NM_001288739.1:<br>c.1197-8G>A | 10a | MS, FS,<br>refractor<br>y | 3<br>month<br>s | Profound DD,<br>GMFCS V,<br>CFCS V | Dystonic movement<br>disorder, profound<br>hypotonia, CVI | Asynchrony,<br>electroclinical<br>myoclonic seizures | Diffuse cerebral<br>atrophy |
| Individual 9 | 3 years | Female | NM_001288739.1:<br>c.1202G>A | 10a | No<br>clinical<br>seizures | 6<br>month<br>s | Profound DD,<br>GMFCS V,<br>CFCS V,<br>MiniMACS V | Profound<br>hypotonia, CVI,<br>nystagmus | Hx of HA, mild<br>background slowing,<br>focal spikes and<br>polyspikes | Abnormal diffusion<br>restriction<br>attributed to<br>vigabatrin toxicity,<br>delayed<br>myelination, mild<br>enlargement of<br>ventricles, small CC |
| Individual 10 | Deceased at 10 months | Male | NM_004408.3:<br>c.1195A>T | 10a | ES, FS,<br>refractor<br>y | 3<br>month<br>s | Profound DD,<br>GMFCS V,<br>CFCS V | Hyperkinetic<br>dystonic movement<br>disorder, profound<br>hypotonia, CVI | HA | Not available |
| Individual 11 | 5 years | Female | NM_004408.3:<br>c.1214C>T | 10b | No<br>seizures | NA | Mild DD,<br>GMFCS I,<br>CFCS I, MACS<br>II | Mild hypotonia,<br>behavioral problems | Moderate increase in<br>beta frequencies | Normal @ 5 years |

ES = epileptic spasms. MS = myoclonic seizures. SS = subclinical seizures. GE= generalized epilepsy. FE= focal epilepsy. CE = combined epilepsy. LGS=Lennox-Gastaut Syndrome. DD= developmental delay. GMFCS = Gross Motor Function Classification System. CFCS = Communication function classification system. CVI = cortical visual impairment. HA = hypsarrhythmia. WM = white matter. SF=seizure free

## Discussion

Here, we report the genetic and clinical features of eleven individuals with *DNM1*-related disorders, including eight individuals with a recurrent *de novo* splicing variant in *DNM1* and three individuals with novel *de novo* variant. We provide several lines of evidence suggesting the identified recurrent *de novo* splicing variant acts through a dominant-negative mechanism, including (1) the syndrome phenocopies the DEE previously ascribed to the established *DNM1*-related disorder typically due to dominant-negative missense variants; (2) functional studies demonstrate the mutant transcript creates a new splice site upstream of the canonical splice site, leading to the insertion of two amino acid residues; (3) structural modeling shows that mutant dynamin-1 forms an oligomeric complex without the capacity for oligomerization-induced GTPase activation; and (4) neuropathological samples reveal an accumulation of synaptic vesicles at the plasma membrane suggestive of impaired vesicular fission. We further demonstrate that variants involving exon 10a lead to a more severe disease than variants affecting exon 10b, likely due to relatively decreased expression of transcripts including exon 10b in the brain.

Among the individuals with variants affecting exon 10a, eight had a recurrent intronic variant, yet their clinical presentations were highly similar to individuals with missense variants expected to result in dominant-negative effects.^17^ This observation is unexpected given that intronic variants near splice sites are typically expected to result in loss-of-function, causing haploinsufficiency as approximately 50% of the protein product is nonfunctional. Instead, we provide evidence here that this splicing variant instead leads to dominant-negative effects, which would recapitulate typical *DNM1*-related disorders at the genotype level.^26^ We predict that this effect is unique to this particular splice site variant, as other splice-site variants, likely leading to loss of function of *DNM1,* are present in the population. For example, NM_004408.3:c.589+1G>C, NM_004408.3:c.1557+1G>T, and NM_001288739.1:c.1335+1G>A are variants located directly within splice donors within the GTPase and middle domain portions of the sequence. All are present in the gnomAD database in apparently healthy individuals and are expected to disrupt splicing, leading to loss-of-function and haploinsufficiency. While intronic or splice site variants leading to dominant negative disease is unusual, it is not unprecedented in genetic developmental and epileptic encephalopathies. A dominant-negative intronic variant has been reported in *GABRG2* (MIM: 137164), as the truncated protein product forms aberrant multimers with other GABA_A_ receptor subunit that fail to traffic to the plasma membrane.^44^ In contrast to the previously identified *GABRG2* variant, we found that this recurrent *DNM1* variant is not protein-truncating, but instead forms an alternate splice acceptor upstream of the normal exon 10a acceptor. The resulting insertion of six bases to the exon and two amino acids to the dynamin-1 protein would be expected to disrupt normal oligomerization, leading to stable multimers expected to lack oligomerization-dependent protein function.

We traced the pathogenic mechanism of this splice variant further with neuropathological specimens from one deceased individual. The dense packing of vesicles directly abutting the plasma membrane **(Figure 6)** demonstrates that vesicles can form from the plasma membrane but are unable to be subsequently removed from the membrane, which is thought to be secondary to impaired vesicular fission. It is possible that the observed synaptic hyperplasia with increased density of synaptic boutons is due to increased axonal sprouting in an effort to circumvent dysfunctional synapses. Similar observations have been made for known dominant-negative missense variants in *DNM1*, with a reduced quantity of cytoplasmic vesicles in mouse models and an excess of membrane invaginations and membrane-adjacent vesicles in HeLa cells.^23^ Thus, our findings affirm that the splicing variant NM_001288739.1:c.1197-8G>A has dominant-negative effects identical to those of known disease-causing missense variants in *DNM1*, consistent with the disease presentation of the eight individuals with this variant, which is indistinguishable from typical *DNM1*-related disorders and inconsistent with carriers of a single loss-of-function allele.

With this affirmation that the recurrent variant operates by a mechanism similar to established pathogenic variants, we were able to classify it as a variant that uniquely affects exon 10a of *DNM1.* Accordingly, taking into account this variant along with the three novel reported disease-causing variants, we find that in spite of the inclusion of exon 10b in the canonical transcript of *DNM1,* exon 10a represents the overwhelmingly dominant cortical isoform in developing brains, accounting for double the proportion of dynamin-1 in cortical samples compared to exon 10b. As a result, dominant-negative missense variants affecting both types or DNM1a alone are expected to result in similar, severe clinical features, as the majority of dynamin-1 proteins are altered in either case. In contrast, variants affecting DNM1b alone would leave the majority of dynamin-1 intact, resulting in a milder expected presentation. The clinical patterns we report confirm these predictions, as all ten individuals with DNM1a-only variants replicated the severe *DNM1* phenotype while the individual with a DNM1b-only variant had significantly milder symptoms.

Exon-dependent variability in disease has already been established in mouse models of *DNM1.* The fitful mouse, a line which was found to have the spontaneous mutation p.Ala408Thr in exon 10a of the mouse homolog *Dnm1*, presents with severe seizure disorder. The fitful phenotype is present in mice with variation in Dnm1a but not Dnm1b, and the heterozygous condition in humans is phenotypically equivalent to the homozygous fitful mouse.^22^ While it was previously thought that this was due to splicing patterns not relevant to human disease, our findings suggest that the need for homozygosity in mouse models reflects the significant skew towards exon 10a in human cortex. In fact, two human families with homozygous nonsense variants in *DNM1* have disease similar to typical *DNM1*-related disorders.^45^ Thus, a vast majority of *DNM1* product in humans and mice must be affected in order to replicate the effects of a single missense variant in human DNM1a, emphasizing the significance of this isoform alone in causing severe disease.

Disease-causing variation in alternative, non-canonical transcripts are also known to cause disease in a wide range of conditions beyond *DNM1.* Pathogenic variants have been identified in alternative coding regions of well-known epilepsy genes such as *CACNA1A* (MIM: 601011) and *STXBP1;* these variants would go undetected if testing were limited to a single, canonical transcript. *CACNA1A* and other voltage-gated calcium channels are particularly prone to alternative patterns of expression which may affect disease phenotypes. For example, a second cistron of *CACNA1A,* which encodes a transcription factor with wider expression than the fulllength channel isoform, has been implicated in some symptoms of spinocerebellar ataxia type 6.^46^ Moreover, we have previously described an individual with a variant in an alternatively spliced exon of *STXBP1* leading to a relatively mild phenotype, including the ability to walk independently, communicate using several words, and no history of seizures.^18^ As such, it is imperative to account for transcript- and isoform-specific clinical patterns in diagnosis and care of individuals with genetic conditions, including *DNM1*-related disorders.

By elucidating transcript-dependent disease-causing variation in *DNM1,* our findings also can inform clinical variant interpretation. Exon 10a of *DNM1* is absent from the Matched Annotation from NCBI and EMBL-EBI (MANE) Select and MANE Plus Clinical transcript sets, which are standardized for use in clinical genetics and variant interpretation.^47^ Future iterations of transcript standardization may benefit from accounting for the DNM1a isoform to facilitate molecular diagnosis in individuals with exon 10a-specific disease-causing variants in *DNM1*. Ultimately, given the evidence that isoforms expressing exon 10a account for the most severe features of *DNM1*-related disorders, our findings illuminate the necessity of assessing the relevant transcript in clinical genetic testing in order to make accurate genetic diagnoses in individuals with possible *DNM1*-related neurodevelopmental disorders.

In summary, we identify DNM1a, characterized by alternative splicing of exon 10a, as the predominant isoform responsible for the classical *DNM1* phenotype in contrast to the milder presentation caused by a variant in DNM1b. Our findings underscore the importance of including the DNM1a sequence in clinical genetic testing, as well as the need more broadly to include relevant transcripts in diagnosis of genetic neurodevelopmental disorders. Finally, we highlight the continued need to consider splice-site variants as causative of dominant-negative disease.

## Supplemental Data

Supplemental data includes one table detailing the cohort from which brain tissue was obtained for bulk RNAseq.

## Acknowledgements

Research of S.D.J.P. and F.C.S. was supported by Conselho Nacional de Desenvolvimento Científico e Tecnológico of Brazil (CNPq) and Rede Mineira de Genômica Populacional e Medicina de Precisão (RED00314-16) of Fundação de Amparo à Pesquisa de Minas Gerais (FAPEMIG). D.B.A. and D.E.V.P. were supported by an Investigator Grant from the National Health and Medical Research Council (NHMRC) of Australia (GNT1174405) and the Victorian Government’s Operational Infrastructure Support Program. The authors acknowledge Azzedine Yacia, Jean Marc Masse, Eve Brochot, Stephanie May, and Vincent Grammont. I.H. was supported by The Hartwell Foundation (Individual Biomedical Research Award), the National Institute for Neurological Disorders and Stroke (K02NS112600, U24NS120854-01, U54NS108874-04), the Eunice Kennedy Shriver National Institute of Child Health and Human Development through the Intellectual and Developmental Disabilities Research Center (IDDRC) at Children’s Hospital of Philadelphia and the University of Pennsylvania (U54HD086984), and by the German Research Foundation (HE5415/3-1, HE5415/5-1, HE5415/6-1, HE5415/7-1). Research reported in this publication was also supported by the National Center for Advancing Translational Sciences of the National Institutes of Health (UL1TR001878), by the Institute for Translational Medicine and Therapeutics’ (ITMAT) at the Perelman School of Medicine of the University of Pennsylvania, and by Children’s Hospital of Philadelphia through the Epilepsy NeuroGenetics Initiative (ENGIN). The project was in part supported by Award number T32NS007413 from the National Institute of Neurological Disorders and Stroke (NINDS). The content is the sole responsibility of the authors and does not necessarily represent the official views of the NINDS of the National Institutes of Health.

## Declaration of Interests

The authors declare no competing interests.

## Web Resources

Online Mendelian Inheritance in Man, http://www.omim.org

## Data and Code Availability

Data are available from the corresponding author on request.

## Notes

### Competing Interest Statement

The authors have declared no competing interest.

